# Lack of consistent effect of dietary fiber on immune checkpoint blockade efficacy across diverse murine tumor models

**DOI:** 10.1101/2025.03.28.645975

**Authors:** Asael Roichman, Gabriela Reyes-Castellanos, Ziqing Chen, Zihong Chen, Sarah J. Mitchell, Michael R. MacArthur, Akshada Sawant, Llewelyn Levett, Jesse Powers, Maria Gomez, Maria Ibrahim, Xincheng Xu, Beianka Tomlinson, Xiang Hang, Yong Wei, Yibin Kang, Eileen White, Joshua D. Rabinowitz

## Abstract

Immune checkpoint blockade (ICB) has transformed cancer treatment, but success rates remain low in most cancers. Recent research suggest that dietary fiber enhances ICB response in melanoma patients and murine preclinical models through microbiome-dependent mechanisms. Yet, the robustness of this effect across cancer types and dietary contexts remains unclear. Specifically, prior literature compared grain-based chow (high fiber) to low-fiber purified diet, but these diets differ also on other dimensions including phytochemicals. Here we investigated, in mice fed grain-based chow or purified diets with differing quantities of isolated fibers (cellulose and inulin), metabolite levels and ICB activity in multiple tumor models. The blood and fecal metabolome were relatively similar between mice fed high- and low-fiber purified diets, but differed massively between mice fed purified diets or chow, identifying the factor as diet type, independent of fiber. Tumor growth studies in three implantable and two spontaneous genetically engineered tumor models revealed that fiber has a weaker impact on ICB (anti-PD-1) efficacy than previously reported. In some models, dietary modulation impacted ICB activity, but not in a consistent direction across models. In none of the models did we observe the pattern expected if fiber controlled ICB efficacy: strong efficacy in both chow and high-fiber purified diet but low efficacy in low-fiber purified diet. Thus, dietary fiber appears to have limited or inconsistent effect on ICB efficacy in mouse models, and other dietary factors that correlate with fiber intake may underlie the clinical correlations between fiber consumption and immunotherapy outcomes.

## Introduction

The development of immune checkpoint blockade (ICB)-based immunotherapy has transformed the field of cancer therapy, giving hope for cure in conditions previously considered untreatable. However, despite this significant advancement, response to ICB therapy varies, ranging from approximately 45-60% in melanoma to only 15-30% in most solid tumors (1,2). Factors associated with favorable response to ICB include DNA mismatch repair deficiency, microsatellite instability, high tumor mutational burden and high PD-L1 expression (3,4). These factors are not, however, under control of the patient. A modifiable factor that is increasingly showing promise to predict responses to ICB in both mice and humans is the gut microbiome (3,4). The gut microbiome - the collection of microorganisms colonizing the colon – plays a major role in anti-tumor immunity, with diverse gut microbiome compositions associated with favorable anticancer response to ICB (5–7). Interventions designed to modify the microbiome into a more favorable composition may therefore have the potential to enhance response rates to ICB therapy.

One way to manipulate microbiome composition is by introducing beneficial bacteria directly through a probiotic cocktail, or by performing fecal-microbiome transplant (FMT) of whole microbial communities from donors with desirable microbial compositions. FMT from ICB-responders has shown promise in some melanoma patients for improving response to ICB, while other patients did not respond favorably (8). One possible reason for treatment failure is that microbes delivered by FMT may not efficiently colonize the recipient’s gut, thereby failing to induce the desired changes in microbiome compositions (8).

An alternative or complementary way to manipulate gut microbiome composition and potentially improve ICB response is through dietary changes (9,10). A key dietary ingredient that feeds the gut microbiome and can be therefore used to influence its composition is dietary fiber (10,11). Dietary fibers are edible carbohydrate polymers that are resistant to endogenous digestion in the small intestine. They can be classified into two categories: (i) insoluble fibers, such as cellulose, hemi cellulose and lignin, which are slowly or minimally digested by the gut bacteria and (ii) soluble fibers, such as inulin, pectin and beta-glucans, which are readily metabolized by the gut microbiota (11). Spencer *et al.* recently investigated the role of dietary fiber in the response to ICB (12). Interestingly, the study found that melanoma patients who consume sufficient fiber (evaluated based on dietary questionnaire) demonstrated improved progression-free survival rate relative to patients with insufficient fiber intake.

As causality could not be inferred from this observational human cohort, results were validated in mice. Specifically, in transplantable melanoma and MC-38 colon carcinoma models, specific pathogen-free (SPF) C57BL/6 mice consuming a grain-based chow, which is high in fiber, showed improved response to ICB (anti-PD-1 or anti-PD-L1) relative to mice fed a diet low in fiber (12). Motivated by these results, clinical trials testing the effect of high-fiber diet on ICB outcomes are ongoing (e.g., NCT04645680, NCT06298734 and NCT04866810).

When conducting dietary intervention studies, it is critical to account for various aspects of the diets. In their mouse studies, Spencer *et al.* investigated the effect of fiber by comparing a grain-based rodent chow as the high-fiber diet (17.6% total fiber) to a purified-ingredient low fiber diet (8% cellulose with no soluble fiber) (12). Notably, grain-based chows and purified diets are inherently different: grain-based chows are manufactured primarily from unrefined ingredients (typically ground corn, soy, wheat and fish), whereas purified diets are prepared from refined, purified ingredients (e.g., casein protein, oils and starch) (13). In terms of fiber, standard grain-based chows typically contain 15-20% fiber of a complex mixture of soluble and insoluble fibers, while purified diets contain specific fibers, typically cellulose (insoluble) and inulin (soluble). As such, chow and purified diets differ not only in fiber composition, but also in multiple other factors including the source of macronutrients, vitamins, minerals, and the presence of phytochemicals (plant secondary metabolites) and heavy metals (13). Any of these factors could potentially contribute to the observed synergy between chow and ICB therapy. To conclusively determine the role of fiber and identify which types may be most effective, it is necessary to compare diets that vary in fiber content but are otherwise identical.

In this study, we explored the effect of dietary fiber on ICB efficacy by feeding mice either typical grain-based chow or purified diets with varying fiber (high/low cellulose/inulin). We used five preclinical tumor models representing varied cancer types and conducted the experiments across different animal facilities. Our key questions were: (i) How significant is the role of fiber in mediating metabolic differences between grain-based chow and purified diets? (ii) Does a grain-based chow consistently enhance response to ICB therapy relative to a low fiber purified diet across tumor models? (iii) Do dietary fibers improve ICB efficacy when tested within a controlled purified diet, and if so, are specific types of fibers, such as insoluble versus soluble fibers, more effective than others? Overall, we find that chow versus purified diet has massive impact on the circulating metabolome largely independent of fiber, and that different diets can impact ICB efficacy in different tumor models, but with no consistent pattern across models and no consistent trend towards high fiber being desirable.

## Results

### Diet design

To study the impact of dietary fiber on response to ICB, we used a grain-based chow diet, which contains a complex fiber mixture (∼ 15.5% w/w insoluble fiber and ∼1.9% w/w soluble), and designed six purified diets varying in their fiber content. The six purified diets closely matched to chow in macronutrient composition (Supplementary Fig. S1A). Two diets were low in fiber and four had high fiber, with cellulose added as insoluble fiber and inulin as soluble fiber. Specifically, the two low-fiber purified diets contained low cellulose (LC, 2.6% w/w) and either no inulin or 0.6% w/w inulin. The four high fiber purified diets contained high cellulose (HC, ∼15.5% w/w, designed to match the amount of insoluble fiber in chow), and variable amounts of inulin (In, 0-8% w/w) (Fig. 1A, see also Supplementary Table S1). The diet with high cellulose and intermediate inulin (1.9%) was designed to contain both insoluble and soluble fiber at similar levels to chow.

**Figure 1.**
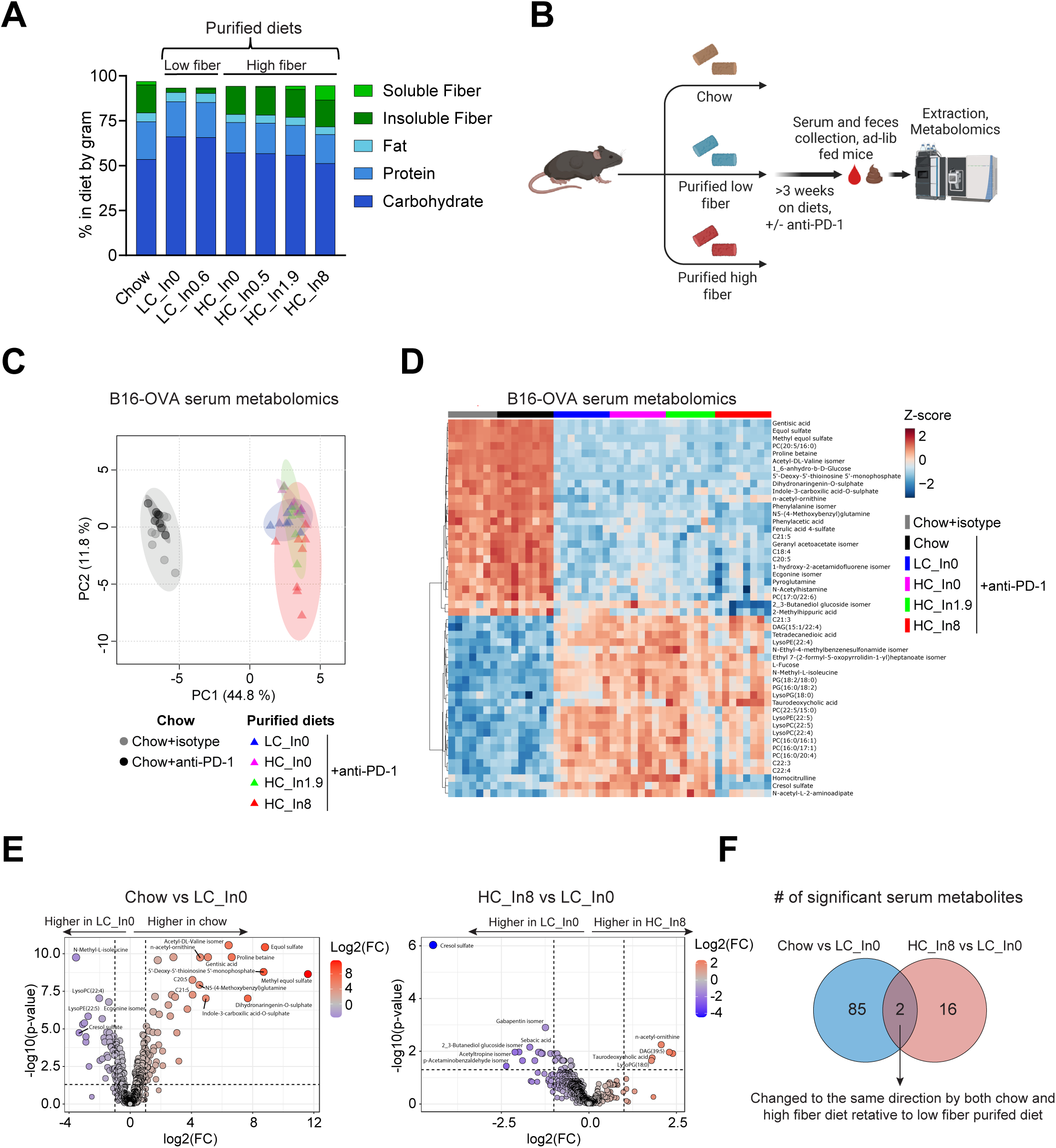
Purified diet has a greater impact than fiber on the serum and fecal metabolome. **A**, Macronutrient and fiber composition of diets used in this study, represented by %gram. In each tumor experiment we used one of the low fiber purified diets and up to four of the high fiber purified diets. LC, Low cellulose (2.5-2.6 %gram). HC, High cellulose (14.9-15.7 %gram). In, Inulin. Numbers in diet names represent %gram of inulin in the diets. **B**, Schematic showing the metabolomics study design. Created with BioRender. **C**, Principal component analysis of serum metabolomics from the B16-OVA experiment. **D**, Heatmap showing log-transformed relative abundances of top 50 significant metabolites by ANOVA of the same metabolomics dataset. **E**, Volcano plots of serum metabolites in mice fed chow vs. low-fiber purified diet (LC_In0) and in mice fed high-fiber (HC_In8) versus low-fiber (LC_In0) purified diet (all B16-OVA tumors and anti-PD-1 treatment). For C-E, *n*=7-8 mice per group. **F**, Venn diagram of significantly changed metabolites from the comparisons shown in panel E, showing minimal overlap between the high-fiber purified diet and chow (only 2 metabolites changing in common versus low-fiber purified diet).

### Grain-based chow and purified diets induce fundamental metabolomic differences independent of fiber content

Diet can influence the response to immune checkpoint blockade (ICB) by altering metabolism (14–16). Thus, we sought to first understand how the addition of fiber to purified diets compares to chow metabolically. To this end, we conducted serum and fecal metabolomics on mice fed either chow or purified diets with low or high fiber (Fig. 1B). To ensure relevance to the anti-PD1 efficacy experiments, metabolomics was conducted in tumor-bearing mice treated with either isotype control or anti-PD1. This was replicated across three tumor models for serum (two for feces) (See Supplementary Tables S2-S4 for full serum metabolomics data). We first assessed the global relationship between all samples in each serum dataset by performing principal component analyses (PCA). In all three datasets, PCA revealed a clear separation along the first principal component (PC1), with a distinct cluster containing chow and another containing all purified diets, while fiber content within purified diets had a minimal impact (Fig. 1C and Supplementary Fig. S1B, C). The clustering was unaffected by whether the mice were treated with ICB (Fig. 1C and Supplementary Fig. S1B, C). These results indicate that diet type, i.e., chow or purified diet, has a far greater impact on driving metabolome changes than ICB or fiber content.

Similarly, in all serum datasets, hierarchical clustering of top significant metabolites was primarily driven by whether the diet was grain-based chow or purified (Fig. 1D and Supplementary Fig. S1D, E). For example, in the B16-OVA model, out of 593 detected metabolites, 87 metabolites significantly changed between chow and low-fiber purified diet (Fold>2, FDR<0.05), whereas only 18 metabolites significantly changed between high-fiber and low-fiber purified diet, with much larger fold changes observed in the chow vs. purified diet comparison (Fig. 1E). Moreover, the metabolite changes induced by chow and high-fiber purified diets showed minimal overlap, with only 2 metabolites exhibiting the same directional change in both chow and high-fiber diets compared to the low-fiber purified diet (Fig. 1F). Similar results were observed in the two other metabolomics datasets (Supplementary Table S5). These findings suggest that the chow-associated circulating metabolome is primarily driven by factors beyond fiber content.

We next analyzed the fecal metabolomics (see full data in Supplementary Tables S6 and S7). More metabolites significantly changed in feces than in serum for both chow vs. purified diet and low-fiber vs. high-fiber purified diet comparisons (Supplementary Table S8). Nevertheless, similar to serum, PCA and hierarchal clustering showed that diet type (i.e., chow vs. purified) had a much larger influence than fiber content in driving fecal metabolite abundances (Supplementary Fig. S1F-I). Collectively, the metabolomics data reveal that grain-based chow fundamentally differs from purified diet in factors that exceed the influence of fiber content.

### Chow, but not by isolated dietary fiber, favors ICB efficacy in the MC-38 colon carcinoma model

We next sought to determine the impact of dietary fiber on ICB efficacy in murine tumor models. We compared standard grain-based chow (high-fiber) to purified diets with either low or high fiber. We were anticipating a pattern where ICB efficacy is strong in chow and in the high-fiber purified diet compared to low-fiber purified diet. We first compared the effects of chow and a low-fiber purified diet (low cellulose, 0.6% inulin) on ICB efficacy in the MC-38 colon carcinoma allograft model in C57BL/6 mice. To assess the consistency across sites, the experiment was conducted both at Charles River Laboratories and Princeton University. Despite somewhat lower calorie intake in mice on the low-fiber purified diet, across both locations, body weight remained comparable among all groups throughout the study (Supplementary Fig. S2A-D). At Charles River, in accordance with Spencer *et al.* (12), chow-feeding slowed tumor progression in isotype control groups and significantly improved ICB response compared to the low-fiber purified diet (Fig. 2B and C). Chow-fed mice also showed a trend for higher rate of complete tumor regression in response to ICB therapy (4/12 for chow vs. 2/12 for low-fiber purified diet, Supplementary Fig. S2E). At Princeton, tumor growth in isotype control groups was comparable between the two diets. However, as seen at Charles River and in Spencer *et al.* (12), chow feeding improved ICB response, though the effect was only marginally significant (Fig. 2D and E), and no complete responders were observed in either group (Supplementary Fig. S2F). These results showed that grain-based chow tends to enhance ICB response in the MC-38 model at both sites.

**Figure 2.**
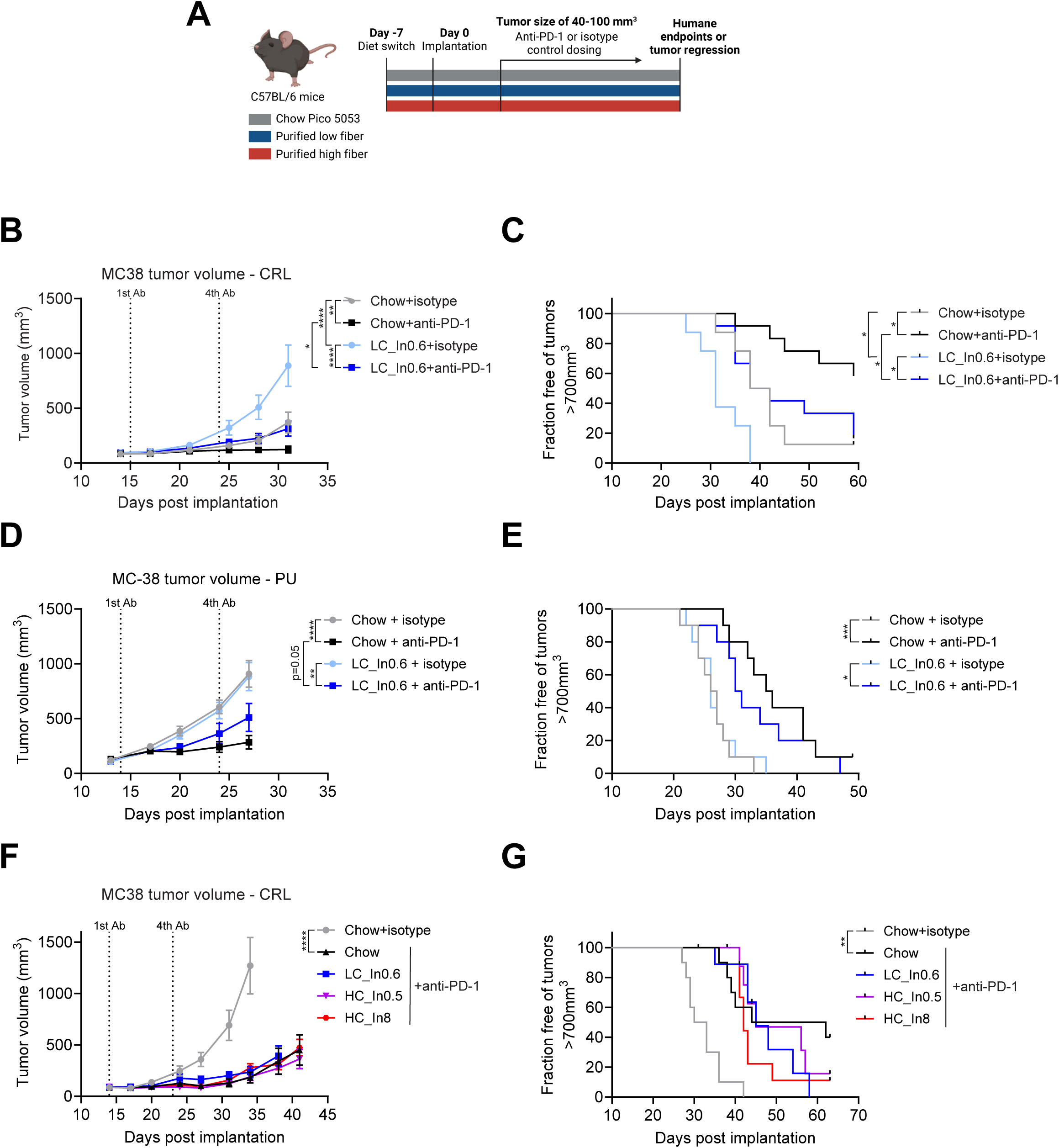
Grain-based chow favors immune-checkpoint blockade efficacy in MC-38 allografts, independently of dietary fiber. **A**, Study design for transplantable tumor models shown in Figures 2 and 3. The first two MC-38 experiments (panels B-E) did not include a purified high-fiber dietary arm. Chow is intrinsically high-fiber. Created with BioRender. **B-C**, Mean tumor volumes and fraction of mice free of tumors larger than 700 mm^3^ in mice implanted with subcutaneous MC-38 tumors, fed chow or a low-fiber purified diet and treated with anti-PD-1 or isotype control, in a study performed in Charles River Laboratories (CRL) facilities. *n*=8 for isotype control and *n*=12 for anti PD-1 groups. **D-E**, Same, in a study performed in Princeton University (PU) facilities. *n*=10. **F-G**, Same, for mice fed chow, or purified diets with low or high insoluble fiber (cellulose, LC or HC, respectively) and low or high soluble fiber (inulin). *n*=10. * *p* < 0.05, ** *p* < 0.01, *** *p* < 0.001, **** *p* < 0.0001, log-rank test for panels C and E and G and two-way ANOVA with Tukey’s post-hoc test at endpoint (day 38 for panel F) for panels B, D and F, values are mean ± SEM.

Next, we aimed to determine how individual fibers in purified diets affect ICB response and how these diets compare to grain-based chow. To this end, we conducted another MC-38 study at Charles River. Mice were fed the same chow and low-fiber purified diet, or two high-fiber purified diets with high cellulose and either low or high inulin. Mice were treated with ICB (and for chow, also isotype control), and body weights and tumor volumes were monitored. Body weights were similar across all groups (Supplementary Fig. S2G). As a positive control, ICB treatment significantly suppressed tumor growth in chow-fed mice compared to isotype control. In this experiment, ICB efficacy in chow did not significantly differ from purified diet (Fig. 2F), although there was a trend towards improved overall survival with chow (fraction free of tumors >700mm^3^, Fig. 2G) and tumor-free survival was observed in the chow group (2/10 for chow vs. 0/10 for low fiber purified diet, Supplementary Fig. S2H). Within purified diets, increasing levels of cellulose or inulin did not affect tumor growth in ICB-treated mice (Fig. 2F and G). Overall, our MC-38 studies suggest that grain-based chow modestly improves antitumor immunity compared to purified diets, while fiber addition does not.

### Purified cellulose, but not by inulin or chow, improves ICB efficacy in one melanoma model, while diet has no effect in another

We next explored the effect of dietary fiber on ICB efficacy in melanoma allograft models. To better capture potential effects of soluble fiber, we used the more extreme version of low soluble fiber diet we designed, switching from 0.6% to zero inulin. We further added high cellulose diets with intermediate inulin (1.9%, matching the soluble fiber content in chow), and high inulin (8%). C57BL/6 mice were fed either chow or one of four purified diets and subcutaneously engrafted with YUMM1.1-9 melanoma cells (17). All groups were treated with ICB, while the chow-fed group also received an isotype control. As a positive control, ICB treatment suppressed tumor growth in chow-fed mice. Notably, there was no significant difference in efficacy between the extreme low-fiber purified diet and chow (Fig. 3A and B). Intriguingly, however, purified diets high in cellulose improved ICB response compared to both chow and the low cellulose purified diet, with the strongest response observed in mice fed the high cellulose/no inulin diet (Fig. 3A and B). These results suggest that insoluble fiber (cellulose), rather than soluble fiber (inulin), enhances ICB efficacy in this model.

**Figure 3.**
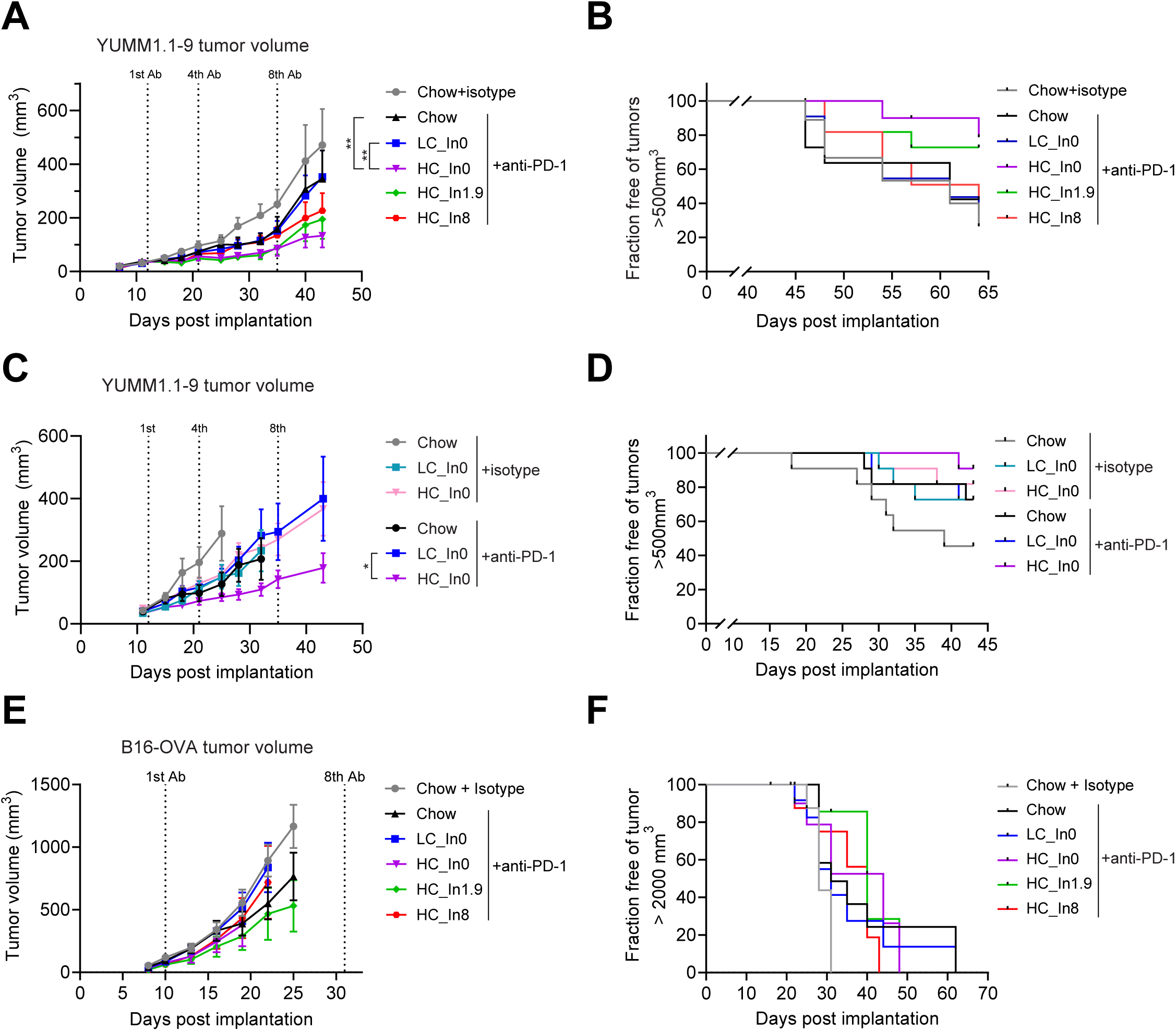
Purified cellulose enhances response to immune-checkpoint blockade therapy in YUMM1.1-9 but not B16-OVA melanoma allografts. **A-B**, Mean tumor volumes and fraction of mice free of tumors larger than 500 mm^3^ for subcutaneous YUMM1.1-9 allografts in mice fed chow or the indicated purified diets and treated with isotype control (for chow) or anti PD-1. *n*=9-11. **C-D**, Similar independent experiment, for mice fed chow, low or high cellulose purified diets and treated with isotype control or anti-PD-1 antibody. *n*=11. **E-F**, Mean tumor volumes and fraction of mice free of tumors larger than 2000 mm^3^ for subcutaneous B16-OVA allografts in mice fed chow or the indicated purified diets and treated with isotype control (for chow) or anti PD-1. *n*=7-12. * *p* < 0.05, ** *p* < 0.01, for panels A, C, and E, by two-way ANOVA with Tukey’s post-hoc test at endpoint, values are mean ± SEM. For panels B, D, and F, log-rank test.

Given the intriguing beneficial effect of insoluble fiber (cellulose) in this model, which appeared to be blunted by adding soluble fiber (inulin), we carried out an additional YUMM1.1-9 study, comparing chow and soluble fiber-free low-cellulose and high-cellulose purified diets. Again in these studies, high-cellulose purified diet resulted in the best tumor control, both in the absence or presence of ICB therapy (Fig. 3C and D). Collectively, these studies suggest that, in the context of YUMM1.1-9 melanoma, purified diet high in insoluble fiber favors tumor control.

Using the same panel of diets in the B16-OVA subcutaneous melanoma model showed no significant differences in ICB efficacy across the diets (Fig. 3E and F), indicating that the impact of fiber on ICB can vary even in different tumors models of the same cancer type. For both melanoma models, food intake was lower in all purified diets relative to chow, with no major effects on body weights (Supplementary Fig. S3A-E). Taken together, these data indicate that cellulose, rather than soluble fiber, improves ICB responses in YUMM1.1-9 melanoma model but not in B16-OVA.

### Cellulose tends to adversely affect ICB efficacy in the Pold1 mutant spontaneous tumor model

Given the results from the YUMM1.1-9 model, we further investigated the effect of cellulose on ICB efficacy using Pold1^D400A/D400A^ mutant mice, a tumor-prone genetic model that we have recently developed that has a proofreading mutation in Polymerase delta1 and as a result has an elevated DNA mutation rate (mutator phenotype) (18). These mice spontaneously develop tumors in various organs, and mostly succumb to thymic and splenic lymphomas before 1 year of age. Prophylactic ICB treatment delays cancer onset and extends median survival in these mice when fed a grain-based chow (18). In contrast to the other mouse cancer models that examine dietary immune tumor control of existing tumors, this spontaneous tumor model also additionally addresses dietary immune impact on tumor initiation. To explore the role of cellulose, male and female Pold1 D400A mice were fed either low cellulose or high cellulose diets (both with no soluble fiber) starting at age of ∼2 months, followed by bi-weekly ICB or isotype control treatments (Fig. 4A). Diet and ICB treatment did not affect body weight or food intake in either sex (Supplementary Fig. S4A-D). Interestingly, ICB treatment delayed early mortality in Pold1 D400A mice that were fed a low cellulose, rather than a high cellulose diet (Fig. 4B, p<0.05 for low cellulose + anti-PD-1 vs. high cellulose + anti-PD-1 for survival up to day 277, which corresponds to the 50th percentile of the total population’s survival). Thus, the effect of cellulose here is opposite to that in YUMM1.1-9 melanoma. In addition, median survival was substantially higher in the low cellulose group treated with ICB (417 vs. 189 days for low cellulose + anti-PD-1 and high cellulose + anti-PD-1, respectively), but these differences did not reach statistical significance due to a greater proportion of late deaths in the low cellulose + anti-PD-1 group (Fig. 4B). These results further indicate that the interaction between dietary fiber and ICB efficacy is cancer-type specific.

**Figure 4.**
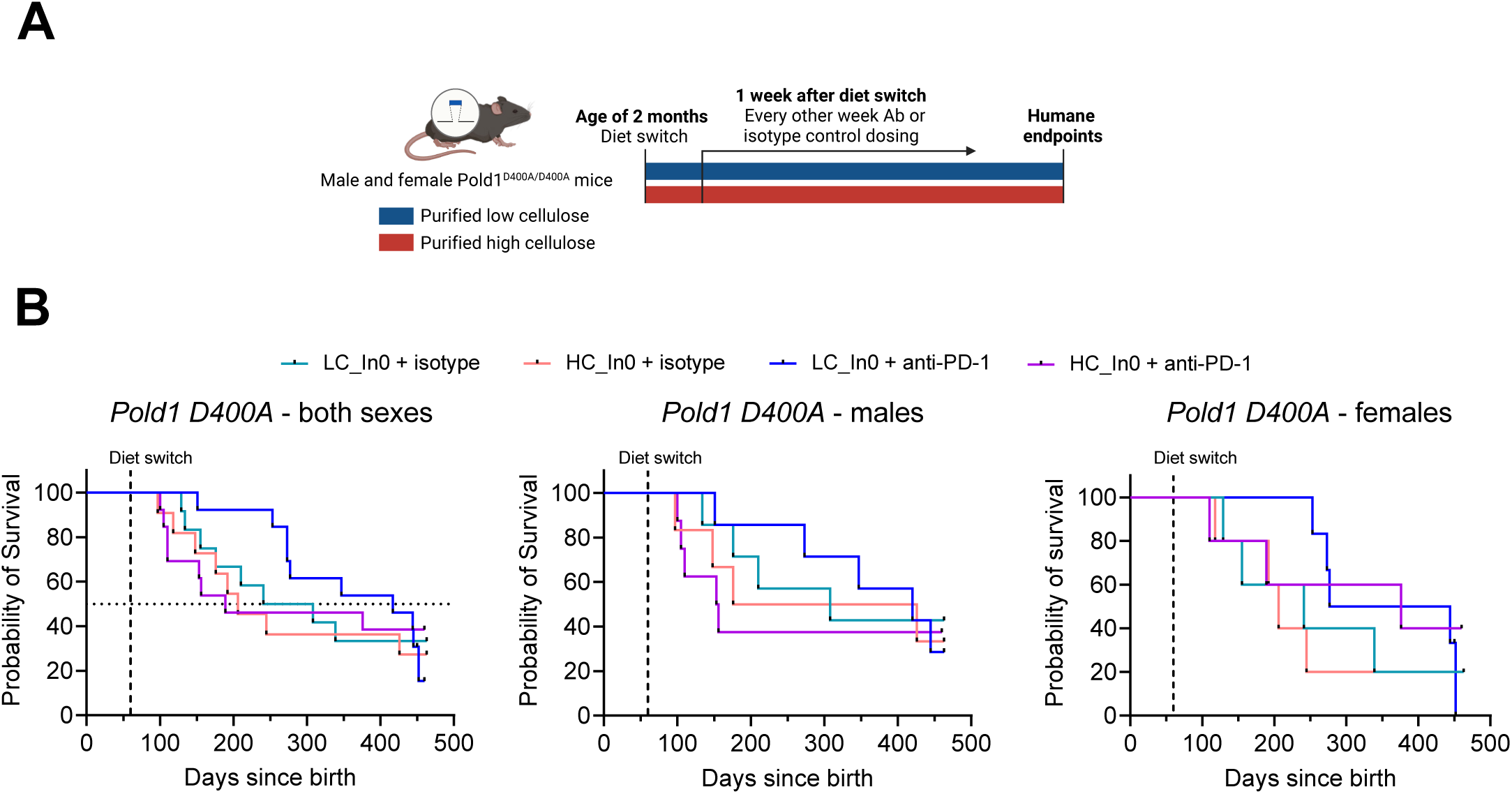
Dietary cellulose tends to negatively impact immune-checkpoint blockade efficacy in Pold1^D400A/D400A^ mice. **A**, Schematic showing Pold1^D400A/D400A^ study design. Created with BioRender. **B**, Survival curves of Pold1^D400A/D400A^ mice fed the indicated high or low cellulose diets and treated with isotype control or anti-PD-1 antibody. Shown are plots for both sexes combined, and males and females separately. *n*=11-13, *n*=6-8, *n*=5-6 mice per group for both sexes, males and females, respectively. Overall survival curves are not significantly different.

### High fiber purified diet, but not chow, modestly improves ICB efficacy in a genetically engineered breast cancer model

We next sought to determine the effect of dietary fiber on ICB efficacy in breast cancer. We used MMTV-PyMT mice, a genetically-induced breast cancer model which develops lung metastases. Mice were fed chow, low fiber purified diet (low cellulose, 0.6% inulin) or high fiber purified diet (high cellulose, 8% inulin) and were untreated or treated with an anti PD-1 antibody. Mice fed high fiber diet gained slightly less body weight over the course of the study, and similar to the transplantable models, mice on purified diets had lower calorie intake (Supplementary Fig. S5A, B). Tumor progression in anti-PD-1 treated mice was slower in mice fed high-fiber purified diet relative to both low-fiber purified diet and chow diets (high-fiber). The anti-tumor effect did not persist to result in a statistically significant survival benefit (Fig. 5A and B). Therefore, the combination of purified diet and dietary fiber modestly improves ICB efficacy in this model.

**Figure 5.**
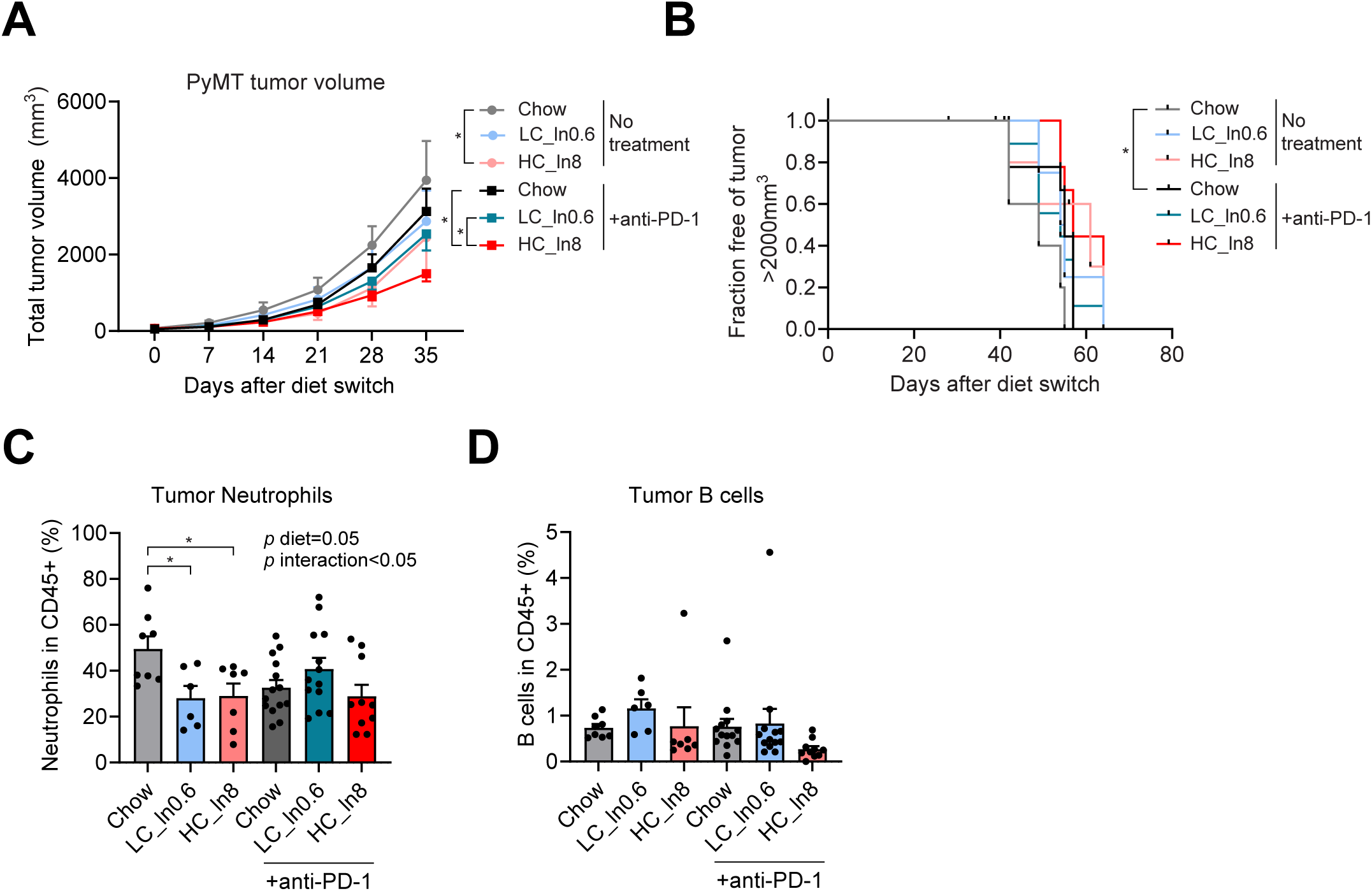
High-fiber purified diet, but not chow, modestly enhances immune-checkpoint blockade efficacy in a breast cancer mouse model. **A**, Mean tumor volumes (calculated as the total volume of all individual mammary tumors per mouse) of MMTV-PyMT mice fed the indicated diets, with or without anti-PD-1 treatment. **B**, Fraction of mice free of a single primary tumor larger than 2000 mm^3^. For A-B, *n*=5 for isotype control and *n*=10 for anti PD-1 groups. **C-D**, Percentage of neutrophils and B-cells among CD45+ cells in primary tumors collected at endpoint. *n*=6-8 tumors collected from 5 mice for no treatment and 10-12 tumors from 9 mice for anti PD-1. *, *p* < 0.05. **, *p* < 0.01. Log-rank test for panel B and ANOVA with Tukey’s post-hoc test for other panels. Values are mean ± SEM.

We next performed immune profiling in tumors collected at ethical endpoint (Supplementary Fig. S6, Supplementary Table S9). Notably, while most immune cell populations measured were not altered, tumor-associated neutrophils, which can contribute to an immunosuppressive environment and promote tumor progression (19), were lower in mice untreated with anti-PD-1 in both purified diets relative to chow (Fig. 5C and Supplementary Fig. S5C). In addition, tumor B-cells tended to be lower in anti-PD-1 treated mice fed high-fiber purified diet (Fig. 5D). Measuring lung metastases nodules at endpoint showed no significant differences between the groups (Supplementary Fig. S5D, E). Thus, in the MMTV-PyMT model, fiber modestly improves ICB response in the context of purified diets, but grain-based chow, despite being rich in fiber, does not show the same effect.

## Discussion

There is a great interest and potential in manipulating diet and/or microbiome to enhance immunotherapy. This is supported by clinical data from multiple trials that show correlations between diet or microbiome composition and ICB outcomes (20,21). Specifically, fiber intake, as deduced from self-reported food diaries, correlated with improved clinical ICB outcomes (12). As fiber consumption likely correlates with many other factors, whether fiber itself drives ICB activity remains to be determined.

Previous mouse experiments compared grain-based chow (high-fiber) to a low-fiber, high-fat purified diet (12). This comparison was confounded by differing macronutrient composition (fat content) and diet purification (which impacts a multitude of aspects beyond fiber including digestibility, phytochemicals and minerals, which trumps fiber in terms of its metabolomic impact). Here we carried out controlled experiments, examining the impact of fiber on ICB activity, finding no consistent role for fiber across 5 different mouse models.

Intriguingly, we did observe consistent dietary effects in particular models. For example, across multiple experiments using MC38 tumors at different sites, chow tended to result in better tumor growth control than low-fiber purified diet, mirroring published results. The challenges were that (i) the benefits of chow were not replicated by adding fiber to the low-fiber purified diet and (ii) trends did not hold across tumor types. For example, diet had no effect on tumor control in the B16-OVA model and high-fiber purified diet (rather than chow) resulted in the best tumor control in the YUMM1.1-9 model. Thus, diet appears to impact immunological tumor control in a context-dependent manner through dietary components beyond fiber.

Our work was limited to murine tumor models and anti-PD1 as the immunotherapy agent. We studied only a single insoluble fiber (cellulose) and soluble fiber (inulin). Different soluble fibers, such as pectins, beta-glucans, and arabinoxylans have distinct effects on the microbiome and its metabolic products (22,23), could easily have differential effects on the immune system, and merit future study.

As opposed to this being a systematic study that applied the exact same dietary conditions across multiple models, experiments were designed sequentially to look for an affirmative effect of fiber (e.g. cellulose was tested in the POLD1 model after it showed positive effects in the YUMM1.1-9 model). Each individual experiment was well-controlled and appropriately powered experiments (each containing substantially more mice than the published affirmative results linking fiber to ICB efficacy), but heterogeneity in experimental designs complicate comparison across the five tumor models including two genetically engineered spontaneous tumor models. The data nevertheless establish the lack of a consistent positive effect in murine ICB of the tested dietary fibers.

This conclusion is important, as it cautions against overly strong endorsement of dietary fiber intake per se for ICB patients until more data is available. Alternative guidance could be to eat a healthy plant-rich diet, as such behavior could benefit patients even if the ultimate causative factor explaining the fiber-ICB correlation ends up being a different plant component. To this end, it would be valuable to re-examine existing clinical data for other potential correlates to ICB efficacy (e.g. plant foods in general; certain fruits, vegetables, grains, or legumes; specific phytochemicals) and for future clinical efforts to take into account dietary complexity beyond fiber.

Patients undergoing immunotherapy are hungry for knowledge of what to eat to maximize their chances of cancer remission. Our results argue for looking beyond the fibers (or at least beyond the commonly studied fibers of cellulose, inulin) and across a breadth of contexts, with the aim of ultimately providing the best possible guidance for future clinical evaluation and patient recommendations.

## Authors’ Contributions

**A. Roichman:** Conceptualization, data curation, formal analysis, writing–original draft. **G, Reyes-Castellanos:** Formal analysis, data curation, validation, investigation. **Ziq. Chen:** Formal analysis, data curation, validation, investigation. **Zih. Chen:** Formal analysis, data curation, validation, investigation. **S.J. Mitchell:** Formal analysis, data curation, validation, investigation. **M.R. MacArthur:** Conceptualization, formal analysis. **A. Sawant:** Investigation. **L. Levett:** Investigation. **J. Powers:** Investigation. **M. Gomez:** Investigation. **M. Ibrahim:** Investigation. **X. Xu:** Investigation. **B. Tomlinson:** Investigation. **X. Hang:** Investigation. **Y. Wei:** Project administration, validation, investigation. **Y. Kang:** Conceptualization, supervision, funding acquisition. **E. White:** Conceptualization, supervision, funding acquisition. **J.D. Rabinowitz:** Conceptualization, supervision, writing–original draft, funding acquisition.

## Acknowledgments

We thank all members of the Rabinowitz, White and Kang labs for their helpful comments on the work. We are also grateful to the staff at the Charles River, Princeton, and Rutgers animal facilities for their dedicated care and support. This work was supported by NIH grant R01 CA243547 and NCI 1OT2CA278609-01, CRUK (CGCATF-2021/100022) awarded to EW, and R01 CA163591 to E.W. and J.D.R., Ludwig Institute for Cancer Research, Ludwig Princeton Branch to E.W., Y.K., and J.D.R., New Jersey Commission on Cancer Research postdoctoral fellowship to A.R. and Ziq.C.

## Methods

### Mice and diets

All mouse studies were performed in accordance with IACUC approved animal protocols at Princeton University (PU), Rutgers Cancer Institute of New Jersey (CINJ) and at Charles River Laboratories (CRL). C57BL/6 mice were purchased from CRL for MC-38 experiment (females), or from Jackson Laboratory (Bar Harbor ME) (#000664) for B16-OVA (females) and YUMM1.1-9 (males) experiments and were at the age of 6-10-week-old at the date of diet switch. Female FVB/N-Tg(MMTV-PyVT)634Mul/J (#002374) mice were purchased from Jackson Laboratory at 6-7 weeks of age. After one-two weeks of acclimation to the facility, mice were switched to one of three experimental diets (Supplementary Table S1). During the acclimation period, mice were fed chow (PicoLab Rodent Diet 5053, LabDiet).

The DNA polymerase delta 1 (*Pold1*) mouse model was created at the Rutgers Cancer Institute together with the Genome Editing Shared Resource. Using CRISPR-Cas9 technology, a point mutation (D400A) was created in the proofreading domain of DNA polymerase δ as described (18). Mice on a C57BL/6 background were used to generate the *Pold1* mouse model. Briefly, *Pold1*^+/+^ mice were crossed with *Pold1^+/D400A^* to obtain heterozygous mice. Then, *Pold1^+/D400A^* × *Pold1^+/D400A^* breeding was used to obtain *Pold1^D400A/D400A^* homozygous mutants. *Pold1^D400A/D400A^* homozygous mice were bred to get experimental mice. The progeny from homozygous breeding was not used for breeding to avoid accumulation of mutations over the generations. PCR amplification and DNA sequencing were used to verify the mutant genotype and only homozygous mice were used in this experiment. At an average of 2 months old, mice were switched from chow diet (PicoLab Rodent Diet 5058) to the LC_In0 or HC_In0 diets.

Body weight and food intake were monitored once or twice per week. During all points of the studies, mice had free access to food and water ad libitum, except PyMT experiment where the mice were overnight fasted before the day of diet switch. Mice were maintained on a 12h light dark cycle.

### Tumor inoculation and endpoints

For transplantable models, one week prior to inoculation, mice were switched to one of the experimental diets (Supplementary Table S1). On day 7, mice were injected subcutaneously with 2.5×10^5^ B16-OVA cells (generously provided by Dr. Marcia Haigis and Dr. Arlene Sharpe), 5×10^5^ MC-38 cells (generously provided by Dr. Arlene Sharpe), or 1×10^6^ Yumm1.1-9 cells resuspended in 0.1 mL PBS. Yumm1.1 cells were obtained from the Bosenberg lab (Yale University, New Haven, CT), and UV irradiated to generate Yumm1.1-9 cells as described (17). Mice were followed twice a week with body weight and tumor volume measurements. Mice were euthanized by CO_2_ asphyxiation if they reached the endpoints: bodyweight loss of >20% of peak bodyweight, body condition score of <=1.5, inappetence/dehydration, tumor diameter >2.0cm, tumor volume >2.0cm^3^, tumor burden >10% of bodyweight, ulceration of tumor or self-mutilation, inability to access food or water, or breathing problems/cyanosis. Tumor volume (mm^3^) was calculated by (length x width x height) x 0.5 [MC-38 in PU, B16-OVA] or by (length x width^2) x 0.5 [MC-38 in CRL, YUMM1.1-9 and PyMT] using digital calipers (Fisher Scientific). Lung nodules (PyMT model) were counted directly after fixation in Bouin’s solution (HT101128, Sigma). The lung images were taken by Zesis SteREO Discovery microscope. For *Pold1* model, necropsy with tissue sampling was performed to confirm the presence of cancer when the mice reached ethical endpoint.

### Anti-PD-1 or isotype control treatment

For transplantable models, mice were randomized and assigned to treatment groups when the average tumor volume reached 40-100 mm^3^. For MC-38 at CRL, mice were treated intraperitoneally with anti-PD-1 (ICH1132, clone RMP1-14, ichorbio) or isotype control (BE0089, Rat IgG2a, BioXcell) at 5 mg/kg twice per week for two weeks. For MC-38 at PU and B16-OVA, mice were dosed with 100ug of anti-PD-1 (BE0273, clone 29F.1A12, BioXcell) or isotype control (BE0089, Rat IgG2a, BioXcell). This was administered as a 100 µl intraperitoneal injection, diluted in *InVivoPure* pH 7.0 Dilution Buffer (IP0070, BioXcell) for Anti-PD-1 or *InVivoPure* pH 6.5 Dilution Buffer (IP0065, BioXcell) for the IgG2a isotype control. Mice were dosed twice per week for two weeks for MC-38, and every 3 days for a total of 8 doses for B16-OVA. For YUMM1.1-9, mice were treated with anti PD-1 antibody (ICH1132, clone RMP1-14, ichorbio) or isotype control (ICH2244, Rat IgG2a, Ichorbio) at 10 mg/kg (both diluted in PBS) every 3 days for a total of 8 doses. For PyMT, mice were randomized and diets were switched at 9 weeks of age, and anti-PD-1 treatment was given at weeks 10-15, twice per week (week 10-12, clone: BE0146, clone RMP1-14, BioXcell; week 13-15, BE0273, clone 29F.1A12, BioXcell). For *Pold1* model, one week after changing diets, mice were randomly assigned to anti PD-1 or isotype groups and treatments were initiated and administered every other week. Mice were injected intraperitoneally with 250 µg/mouse of anti PD-1 antibody (BE0146, clone RMP1-14, BioXcell) or isotype (BE0089, Rat IgG2a, BioXcell) diluted in 100 µl of PBS.

### Serum & feces sampling

Blood was collected by tail snip in ad lib fed mice: For B16-OVA around 9am, the day after the 6^th^ dose of anti-PD-1 or isotype control, for YUMM1.1-9, at 6-8 am, the day after the 3^rd^ dose of anti-PD-1 or isotype control, for PyMT, at 2-3 pm the day after the 5^th^ dose of anti-PD-1 or isotype control. Blood was centrifuged at 10,000 g, 4°C for 10 min. Serum was isolated from the supernatant and stored at -80°C until further analysis. Feces samples were collected at the same time during blood sampling and stored at -80°C until further analysis.

### Metabolite extraction

For serum samples from B16-OVA, YUMM1.1-9, and PyMT studies, 2.5 µL of serum was added into 80 µl of ice-cold methanol and stored in -80°C for at least 30min. Samples were centrifuged at max speed (21,380 g), 4°C for 20 min. Supernatants were collected for LC-MS analysis. For feces samples from B16-OVA and YUMM1.1-9 studies, frozen feces samples were transferred to Eppendorf tubes with ceramic beads on dry ice and disrupted by cryomill (Retsch). About 10 mg of homogenized powder was weighed and extracted by ice-cold acetonitrile: methanol: water (40:40:20, supplemented with 0.5 % vol formic acid). Extracts were vortexed for 10 s, kept on dry ice for 10 min, and neutralized by NH4HCO3 (15% in water, 8.8% vol/vol of extraction buffer was used). Neutralized extracts were vortexed for 10 s again, kept on dry ice for 1h, and centrifuged at 21, 380 g, 4°C for 20 min. Supernatants were collected for LC-MS analysis.

### LC-MS method

Metabolites were separated by hydrophilic interaction liquid chromatography (HILIC) with an XBridge BEH Amide column (2.1 mm × 150 mm, 2.5 μm particle size; Waters, 196006724). The column temperature was set at 25°C. Solvent A was 95 vol% H2O 5 vol% acetonitrile (with 20 mM ammonium acetate, 20 mM ammonium hydroxide, pH 9.4). Solvent B was acetonitrile. Flow rate was 0.15 mL/min. The LC gradient was: 0-2min, 90% B; 3-7min, 75% B; 8-9 min, 70% B; 10-12 min, 50% B; 13-14 min, 25% B; 16-20.5 min, 0.5% B; 21-25 min, 90%. MS analysis was performed on Thermo Fisher’s Q Exactive Plus (QE+) Hybrid Quadrupole-Orbitrap, Orbitrap Exploris 240, or Orbitrap Exploris 480 mass spectrometer, with injection volume of 5-10µl. Two scans were performed in negative mode (m/z of 70-1000) and positive mode (m/z of 119-1000), respectively. Other parameters are listed in the table below:

| MS | Polarity | m/z range | Resolution | AGC target | Max IT |
| --- | --- | --- | --- | --- | --- |
| QE+ | Negative | 70-1000 | 140, 000 | 3e6 | 300 ms |
| QE+ | Positive | 119-1000 | 70, 000 | 3e6 | 300 ms |
| 240 | Negative | 70-1000 | 120, 000 | 1000% | 500 ms |
| 240 | Positive | 119-1000 | 120, 000 | 500% | 500 ms |
| 480 | Negative | 70-1000 | 120, 000 | 1000% | 500 ms |
| 480 | Positive | 119-1000 | 120, 000 | 1000% | 500 ms |

### Immune profiling sample preparation and flow cytometry

For immune profiling in the PyMT model, 0.5 g primary tumor were harvested and dissected in 1.5 ml tubes prior to digestion with tumor dissociation kit (130-096-730, Miltenyi Biotec). Tumors were then dissociated to single cell suspensions according to the manufacturer’s instruction. Cells were spun down at 500g for 5 mins, supernatant removed and then resuspended in 3 ml ACK buffer for red blood cell lysis. Cells were briefly vortexed, allowed to sit at room temperature for 1-3 mins and then quenched with 10 ml of DPBS. Cells were then spun down and resuspended in FACS buffer (2% FBS contained DPBS) for experiment. Single cell suspensions were first incubated with anti-mouse FcγIII/II (CD16/CD32) receptor blocking antibodies (Thermo) for 10 mins followed by staining with antibodies (Supplementary Table S10) for 30mins at 4 °C. Fixable viability dye Near IR (Thermo) or Aqua (Thermo) were used to exclude dead cells. All data were collected on an Attune NxT Flow cytometer and analyzed with FlowJo 10.7.1. Gating for experiments is shown in Supplementary Fig. S6.

### Data and statistical analysis

Raw data was collated in Microsoft excel before being transferred to GraphPad Prism for analysis. Metabolomics data was analyzed using MetaboAnalyst 5.0 (https://www.metaboanalyst.ca/). Data are presented as mean ± SEM for bar and line graphs. For multiple-group comparisons we used one-way or two-way analysis of variance (ANOVA) with Sidak’s or Tukey’s multiple comparison correction. Survival was analyzed by Kaplan–Meier survival curves, p-values were calculated by log-rank or Wilcoxon test using GraphPad Prism and corrected for multiple comparisons by Benjamini-Hochberg method. All comparisons were two-tailed, and a p-value below 0.05 was considered a statistically significant difference. Significant outliers determined by Grubb’s test were removed from final analysis.

## Data availability

All raw data generated in this study are available upon request from the corresponding authors.

---

**Supplementary Figure 1.**
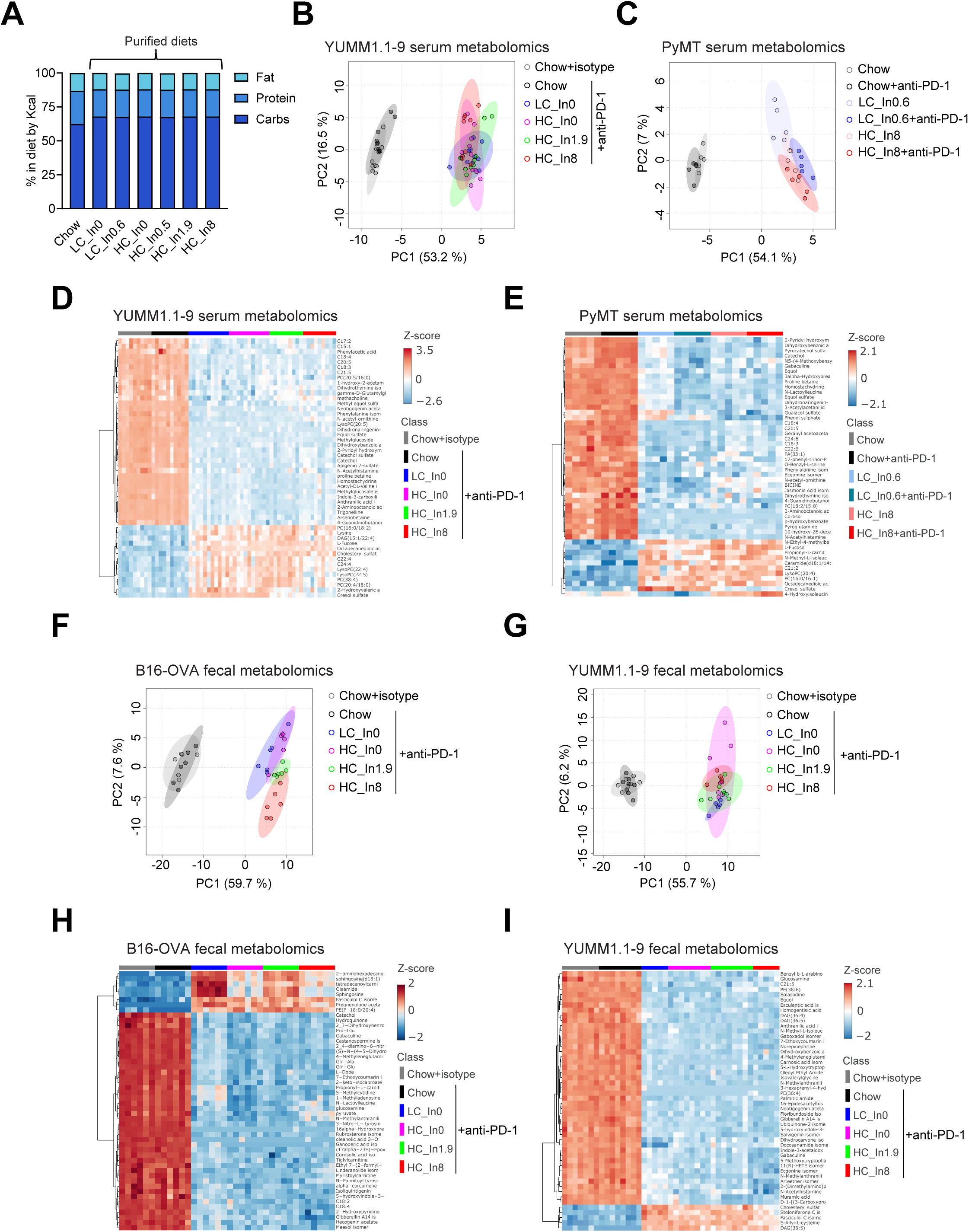
**A**, Macronutrient composition of diets used in this study, represented by %Kcal. **B-C**, Principal component analysis of serum metabolomics from YUMM1.1-9 and PyMT studies. **D-E**, Heatmap showing log-transformed relative abundances of top 50 metabolites by ANOVA from the YUMM1.1-9 and PyMT studies. For B-E, *n*=9-11 mice per group for YUMM1.1-9 and *n*=5 mice per group for PyMT. **F-G**, Principal component analysis of fecal metabolomics from the B16-OVA and YUMM1.1-9 studies. **H-I**, Heatmap showing log-transformed relative abundances of top 50 metabolites by ANOVA from the B16-OVA and YUMM1.1-9 studies. For F-I, *n*=6 mice per group for B16-OVA and *n*=5-8 mice per group for YUMM1.1-9

**Supplementary Figure 2.**
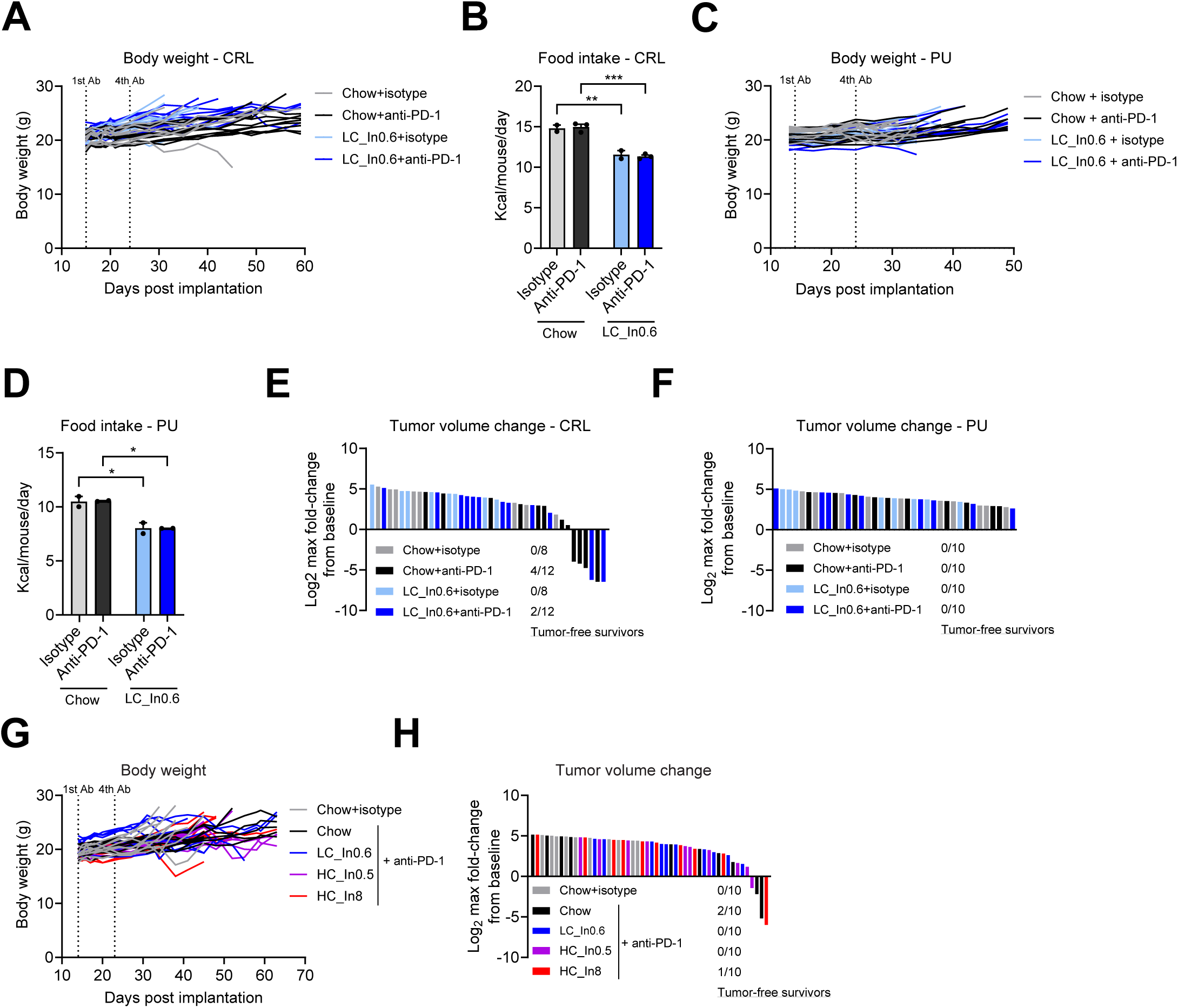
**A-B**, Body weights of individual mice and food intake averaged from 2-3 cages per group for the MC-38 experiment performed at Charles River Laboratories (CRL). Values in B are mean ±SEM. *n* = 8 for isotype control and *n* = 12 for anti PD-1 groups. **C-D**, Same, for the MC-38 experiment performed at Princeton University (PU). *n* = 10, food intake averaged from 2-3 cages per group. Values in D are mean ±SEM. **E**, Maximal tumor volume change from baseline, with the number of tumor-free survivors for each group indicated, in the experiment performed at CRL. *n* = 8 for isotype control and *n* = 12 for anti PD-1 groups. Panels A, B and E correspond to the study presented in main text Figures 2B and C. **F**, Same, for the experiment performed at PU. *n* = 10. Panels C, D and F correspond to the study presented in main text Figures 2D and E. **G-H**, Body weights of individual mice and maximal tumor volume change from baseline with tumor free survivors indicated, corresponding to the study presented in Figures 2F and G of the main text. *n* = 10.

**Supplementary Figure 3.**
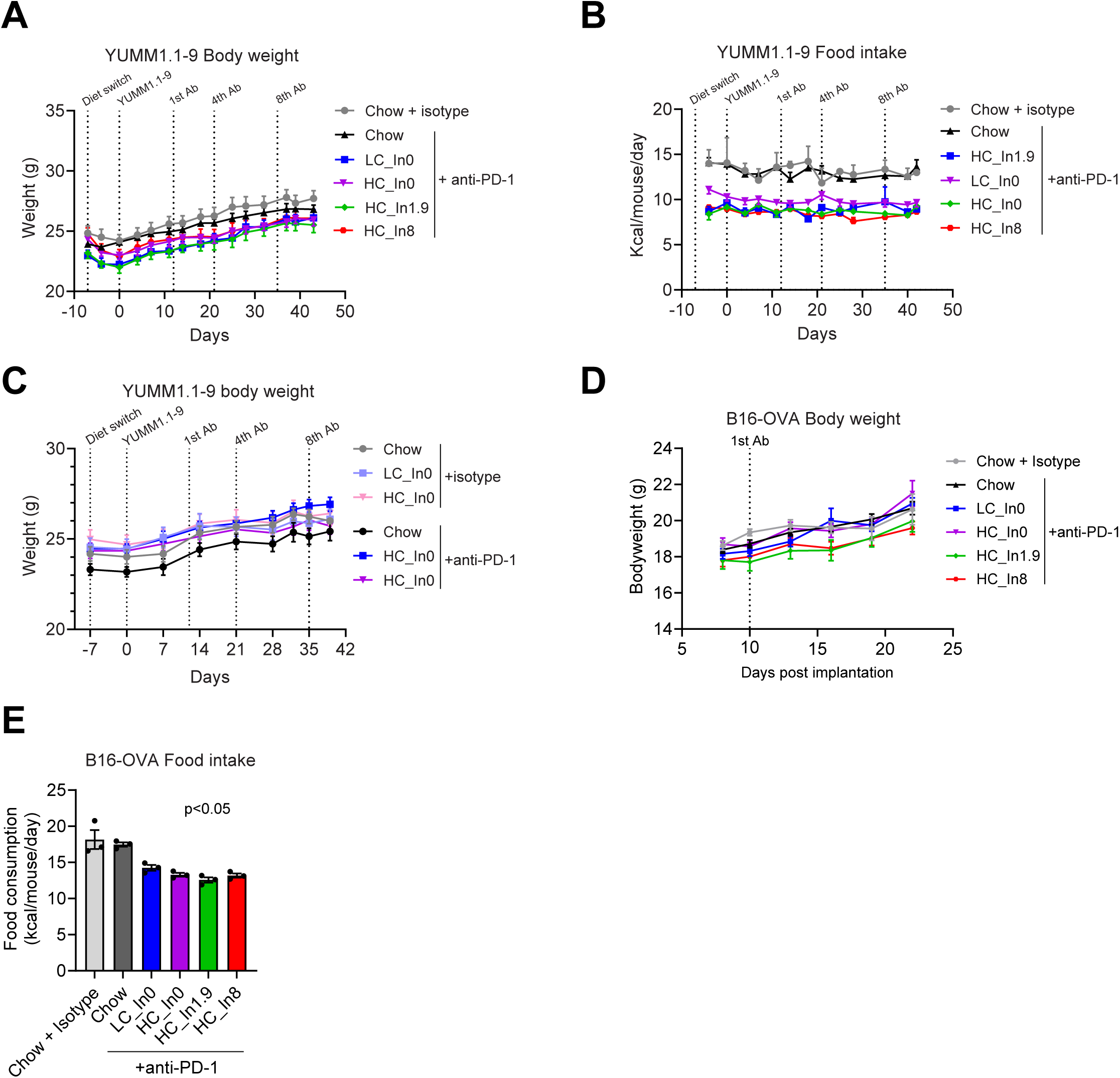
**A-B**, Mean body weight and food intake of mice bearing subcutaneous YUMM1.1-9 allografts from the experiment shown in Figures 3A and 3B of the main text. *n* = 9-11 for A, *n* = 2-3 cages for B. *p*<0.05 for A and *p*<0.0001 for B by two-way ANOVA for main diet effect. **C**, Mean body weights of mice bearing subcutaneous YUMM1.1-9 allografts from the experiment presented in main text figures 3C and D. *n* = 11. **D-E**, Mean body weight and food intake of mice bearing subcutaneous B16-OVA allografts from the experiment presented in main text figures 3E and F. *n* = 7-12 for D, *n* = 3 cages for E. *p*<0.05 for E by one-way ANOVA. For all panels, error bars indicate SEMs.

**Supplementary Figure 4.**
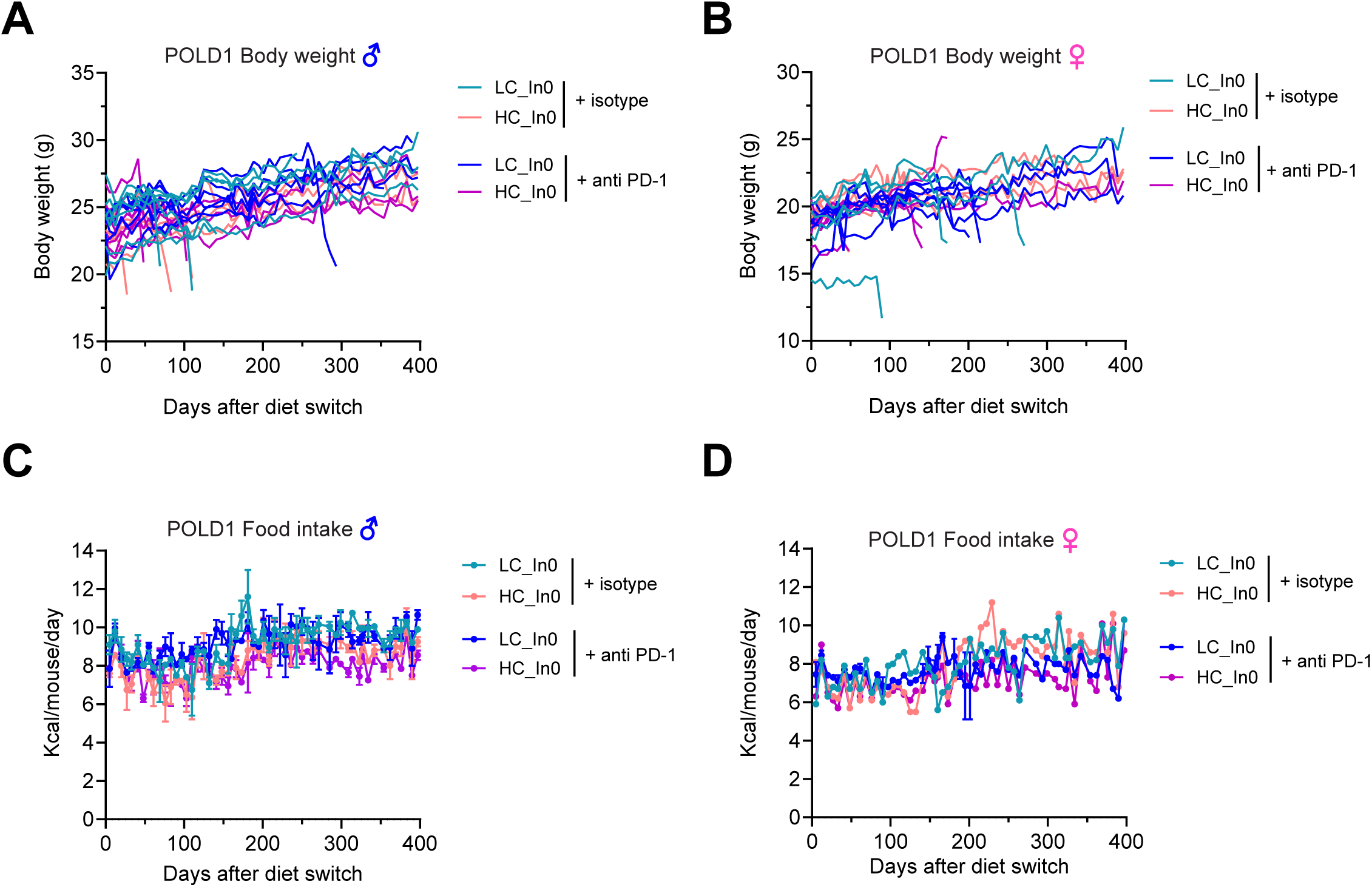
**A-B**, Body weights of Pold1^D400A/D400A^ male and female mice receiving high or low cellulose diet and treated with anti-PD-1 or isotype control antibody. *n*=6-8 for males, *n*=5-6 for females. **C-D**, Food intake of the same mice. *n*=2 cages for males, *n*=1-2 cages for females. Values are mean ± SEM.

**Supplementary Figure 5.**
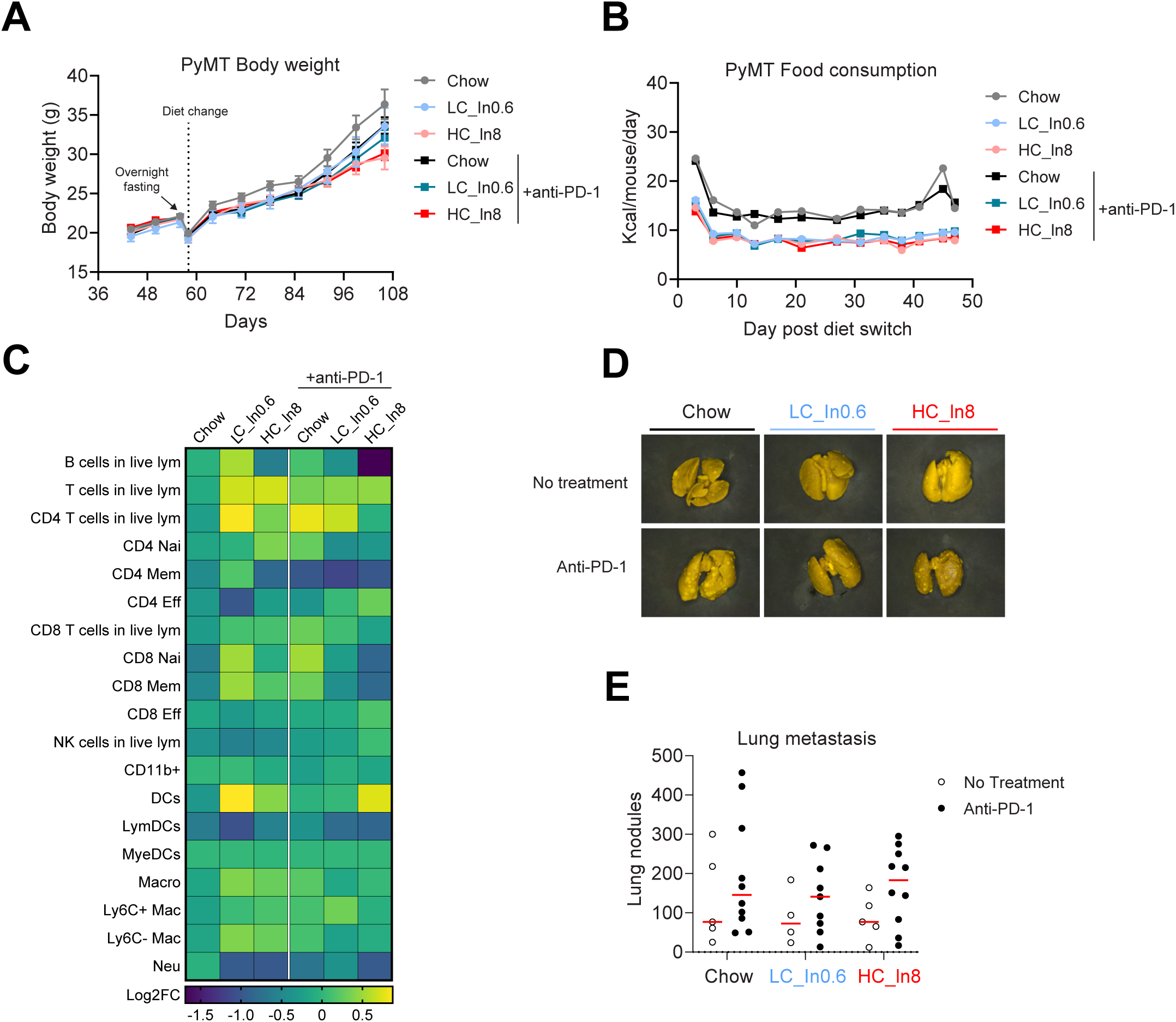
**A-B**, Mean body weight (± SEM) and mean food intake of MMTV-PyMT mice from the experiment presented in main text figure 5A and B. *n* = 5 for isotype control and *n* = 10 for anti PD-1 groups for panel A, *n* = 1-2 cages for panel B. **C,** Heatmap showing the percentage of immune cell populations in primary tumors relative to the chow no-treatment group. Each square represents the average of *n*=6-8 tumors collected from 5 mice for no treatment and 10-12 tumors from 9 mice for anti PD-1. See Supplementary Table S9 for raw data. **D-E**, Representative images and quantification of lung metastases at humane endpoints from the experiment presented in main text figure 5A and B. *n* = 4-5 for no treatment and *n* = 9-10 for anti PD-1 groups.

**Supplementary Figure 6.**
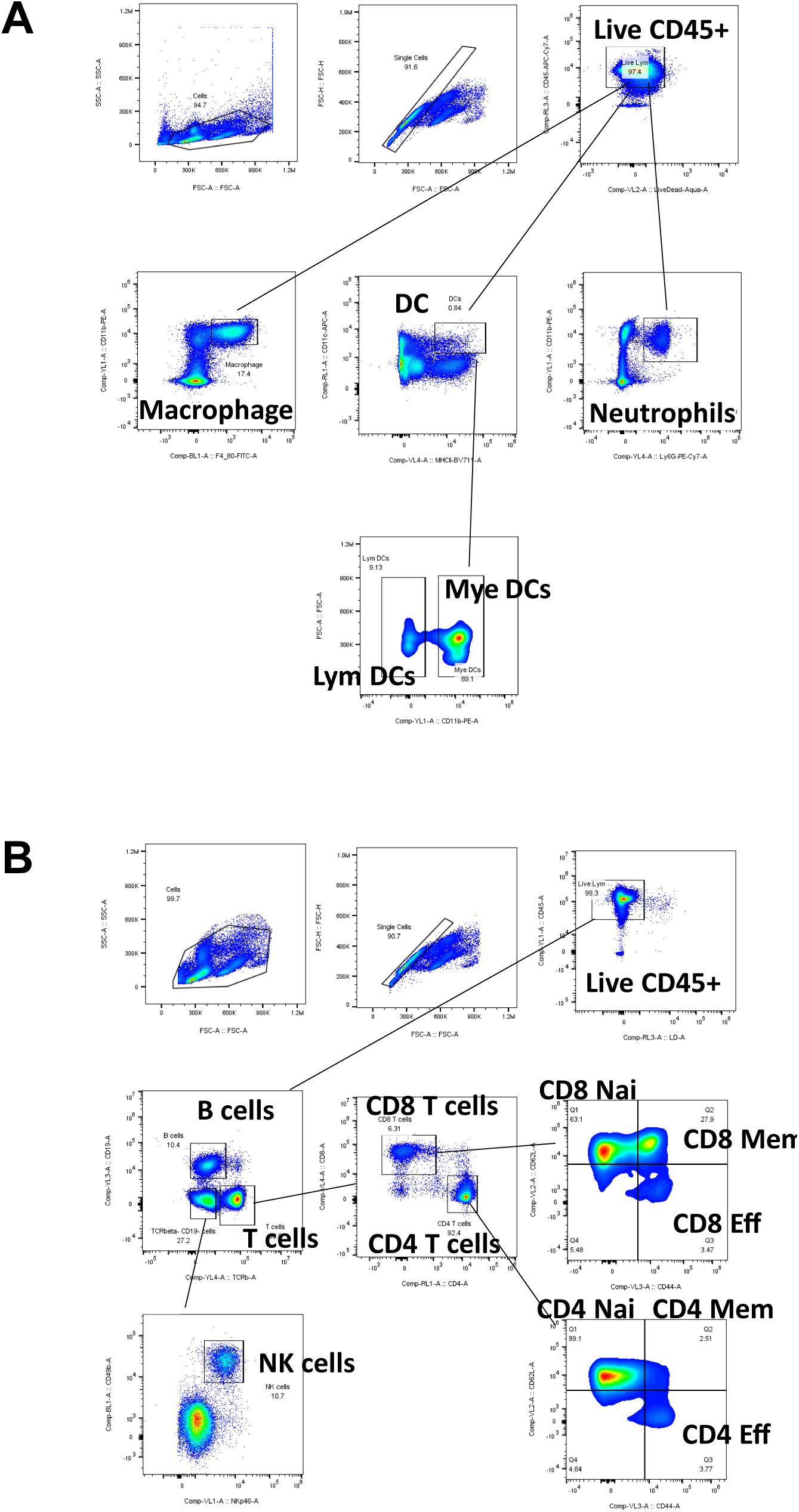
**A**, Gating strategy for myeloid panel. Cells were first gated on singlet and live CD45^+^ live lymphocytes. Then further gated on CD11b+F4/80+ macrophage, CD11b+Ly6G+ neutrophils. CD11c+MHCII+ DCs were further divided into Myeloid derived DCs (CD11b+ DCs) and Lymphoid derived DCs (CD11b-DCs). **B**, Gating strategy for lymphocyte panel. Cells were divided into B (CD19+ TCRb-) and T cells (CD19-TCRb+) after singlet/live CD45+ gate. NK cells were gated on CD19-TCRb-NKp46+CD49b+. T cells were divided into CD4 and CD8 T cells. The memory, effector and naïve state could be identified as CD62L+ CD44+, CD62L-CD44+, CD62L+CD44-respectively. See Supplementary Table S10 for list of antibodies used.

## References

1. Sharma P, Hu-Lieskovan S, Wargo JA, Ribas A. Primary, Adaptive, and Acquired Resistance to Cancer Immunotherapy. Cell [Internet]. 2017;168(4):707–23. Available from: 10.1016/j.cell.2017.01.017

2. Das S, Johnson DB. Immune-related adverse events and anti-tumor efficacy of immune checkpoint inhibitors. J Immunother Cancer. 2019;7(1):1–11.

3. Yamaguchi H, Hsu JM, Sun L, Wang SC, Hung MC. Advances and prospects of biomarkers for immune checkpoint inhibitors. Cell Reports Med [Internet]. 2024;5(7):101621. Available from: 10.1016/j.xcrm.2024.101621

4. Holder AM, Dedeilia A, Sierra-Davidson K, Cohen S, Liu D, Parikh A, et al. Defining clinically useful biomarkers of immune checkpoint inhibitors in solid tumours. Nat Rev Cancer [Internet]. 2024;24(7):498–512. Available from: 10.1038/s41568-024-00705-7

5. Gopalakrishnan V, Spencer CN, Nezi L, Reuben A, Andrews MC, Karpinets T V., et al. Gut microbiome modulates response to anti-PD-1 immunotherapy in melanoma patients. Science (80-). 2018;359(6371):97–103.

6. Tanoue T, Morita S, Plichta DR, Skelly AN, Suda W, Sugiura Y, et al. A defined commensal consortium elicits CD8 T cells and anti-cancer immunity. Nature [Internet]. 2019 Jan 23;565(7741):600–5. Available from: 10.1038/s41586-019-0878-z

7. Sivan A, Corrales L, Hubert N, Williams JB, Aquino-Michaels K, Earley ZM, et al. Commensal Bifidobacterium promotes antitumor immunity and facilitates anti-PD-L1 efficacy. Science (80-). 2015;350(6264):1084–9.

8. Davar D, Dzutsev AK, McCulloch JA, Rodrigues RR, Chauvin JM, Morrison RM, et al. Fecal microbiota transplant overcomes resistance to anti-PD-1 therapy in melanoma patients. Science (80-). 2021;371(6529):595–602.

9. David LA, Maurice CF, Carmody RN, Gootenberg DB, Button JE, Wolfe BE, et al. Diet rapidly and reproducibly alters the human gut microbiome. Nature [Internet]. 2014 Jan 11;505(7484):559–63. Available from: 10.1038/nature12820

10. Zeng X, Xing X, Gupta M, Keber FC, Lopez JG, Lee Y-CJ, et al. Gut bacterial nutrient preferences quantified in vivo. Cell [Internet]. 2022 Sep;185(18):3441–3456.e19. Available from: https://www.biorxiv.org/content/10.1101/2022.01.25.477736v1%0Ahttps://www.biorxiv.org/content/10.1101/2022.01.25.477736v1.abstract

11. Makki K, Deehan EC, Walter J, Bäckhed F. The Impact of Dietary Fiber on Gut Microbiota in Host Health and Disease. Cell Host Microbe. 2018;23(6):705–15.

12. Spencer CN, McQuade JL, Gopalakrishnan V, McCulloch JA, Vetizou M, Cogdill AP, et al. Dietary fiber and probiotics influence the gut microbiome and melanoma immunotherapy response. Science (80-) [Internet]. 2021 Dec 24;374(6575):1632–40. Available from: https://www.science.org/doi/10.1126/science.aaz7015

13. Pellizzon MA, Ricci MR. The common use of improper control diets in diet-induced metabolic disease research confounds data interpretation: The fiber factor. Nutr Metab. 2018;15(1):1–6.

14. Bader JE, Voss K, Rathmell JC. Targeting Metabolism to Improve the Tumor Microenvironment for Cancer Immunotherapy. Mol Cell [Internet]. 2020;78(6):1019–33. Available from: 10.1016/j.molcel.2020.05.034

15. Jia D, Wang Q, Qi Y, Wang Y, Chen S. Microbial metabolite enhances immunotherapy efficacy by modulating T cell stemness in pan-cancer. Cell [Internet]. 2024;187(7):1651–1665.e21. Available from: 10.1016/j.cell.2024.02.022

16. Joachim L, Göttert S, Sax A, Steiger K, Neuhaus K, Heinrich P, et al. Articles The microbial metabolite desaminotyrosine enhances T-cell priming and cancer immunotherapy with immune checkpoint inhibitors. eBioMedicine [Internet]. 2023;97:104834. Available from: 10.1016/j.ebiom.2023.104834

17. Poillet-Perez L, Sharp DW, Yang Y, Laddha S V., Ibrahim M, Bommareddy PK, et al. Autophagy promotes growth of tumors with high mutational burden by inhibiting a T-cell immune response. Nat Cancer [Internet]. 2020;1(9):923–34. Available from: 10.1038/s43018-020-00110-7

18. Sawant A, Shi F, Lopes EC, Hu Z, Abdelfattah S, Baul J, et al. Immune Checkpoint Blockade Delays Cancer and Extends Survival in Murine DNA Polymerase Mutator Syndromes. bioRxiv [Internet]. 2024; Available from: https://www.biorxiv.org/content/10.1101/2024.06.10.597960v1

19. Faget J, Peters S, Quantin X, Meylan E, Bonnefoy N. Neutrophils in the era of immune checkpoint blockade. J Immunother Cancer. 2021;9(7):1–15.

20. Szczyrek M, Bitkowska P, Chunowski P, Czuchryta P, Krawczyk P, Milanowski J. Diet, Microbiome, and Cancer Immunotherapy—A Comprehensive Review. Nutrients [Internet]. 2021 Jun 28;13(7):2217. Available from: https://library.seg.org/doi/10.1190/1.9781560802662.ch1

21. Simpson RC, Shanahan ER, Scolyer RA, Long G V. Towards modulating the gut microbiota to enhance the efficacy of immune-checkpoint inhibitors. Nat Rev Clin Oncol. 2023;20(October):697–715.

22. Patnode ML, Beller ZW, Han ND, Cheng J, Peters SL, Terrapon N, et al. Interspecies Competition Impacts Targeted Manipulation of Human Gut Bacteria by Fiber-Derived Glycans. Cell [Internet]. 2019;179(1):59–73.e13. Available from: 10.1016/j.cell.2019.08.011

23. Murga-garrido SM, Hong Q, Cross TL, Hutchison ER, Han J, Thomas SP, et al. Gut microbiome variation modulates the effects of dietary fiber on host metabolism. Microbiome. 2021;9(117):1–26.

